# Improved T cell surfaceomics by depleting intracellularly labelled dead cells

**DOI:** 10.1101/2025.09.30.679609

**Authors:** Christofer Daniel Sánchez, Aswath Balakrishnan, Blake Krisko, Bulbul Ahmmed, Luna Witchey, Oceani Valenzuela, Minas Minasyan, Anthony Pak, Haik Mkhikian

**Affiliations:** Department of Pathology and Laboratory Medicine, University of California, Irvine, California, USA

**Author notes:** These authors contributed equally to this work (Christofer Daniel Sánchez, Aswath Balakrishnan). For correspondence: Haik Mkhikian,.

**Keywords:** Surfaceome, plasma membrane, glycosylation, intracellular pool, biotinylation

## Abstract

Although the plasma membrane (PM) is among the most biologically important and therapeutically targeted cellular compartments, it is among the most challenging to faithfully capture using proteomic approaches. The quality of quantitative surfaceomics data depends heavily on the effectiveness of the cell surface enrichment used during sample preparation. Enrichment improves sensitivity for low abundance PM proteins and ensures that the changes detected reflect PM expression changes rather than whole cell changes. Cell surface biotinylation with PM-impermeable, amine-reactive reagents is a facile, accessible, and unbiased approach to enrich PM proteins. For unclear reasons however, it results in unexpectedly high contamination with intracellular proteins, reducing its utility. We report that biotinylating human cells with amine-reactive reagents intracellularly labels a small but reproducible population of non-viable cells. Although these dead cells represent only 5±2% of the total, we find that in T cell preparations the dead cells account for 90% of labelled proteins. Depleting Annexin V positive dead T cells post-labelling removes ∼99% of the intracellularly labelled cells, resulting in markedly improved PM identifications, peptide counts, and iBAQ intensities. Correspondingly, we found substantial depletion of intracellular proteins, particular of nuclear origin. Overall, the cumulative intensity of PM proteins increased from 4% to 55.8% with dead cell depletion. Finally, we demonstrate that immature ER/Golgi glycoforms of CD11a and CD18 are selectively removed by dead-cell depletion. We conclude that high intracellular labelling of non-viable cells is the major source of intracellular protein contaminants in amine-reactive surface enrichment methods and can be reduced by dead-cell depletion post-labelling, improving both sensitivity and accuracy of plasma membrane proteomics.

## Introduction

The cell surface proteome, or surfaceome, serves as a critical interface between a cell and its environment, orchestrating essential functions such as signal transduction, nutrient uptake, and immune recognition (1). Consistent with its biological importance, 50% of small molecule drugs and virtually all immunotherapies target plasma membrane (PM) proteins(2, 3). Most PM proteins are transmembrane glycoproteins synthesized in the endoplasmic reticulum (ER)(4). They traffic through the secretory pathway where their glycans are variably processed, delivering a heterogenous set of glycoforms to the PM(5). After variable PM retention times, these glycoproteins are internalized and further trafficked through the endosomal system, frequently being recycled to the cell surface before finally being degraded in lysosomes(6).

Given the biological importance and therapeutic relevance of the PM compartment, there is a need for techniques that allow accurate and sensitive profiling of the surfaceome. Mass spectrometry-based proteomics approaches are powerful in this regard but must be combined with PM enrichment. Enrichment is required to both increase sensitivity for low abundance PM proteins, which collectively constitute ∼1% of the cellular proteome, and to distinguish cell surface from intracellular pools of PM proteins(7). Several strategies have been developed to enrich PM proteins, including density gradient fractionation, and chemical and enzymatic labelling methods coupled to affinity purification(8). The most prominent direct chemical labelling approaches target amines (lysines) or aldehydes (glycans) with membrane impermeable biotinylation reagents such as sulfo-NHS-biotin(9).

Glycan labelling results in fewer intracellular contaminants than amine labelling, but relies on the presence of glycans. Though unglycosylated PM proteins are in the minority, they escape capture by glycan labelling. Furthermore, sialylated glycans are more readily labelled by these reagents(10). Thus, quantitative comparisons, and particularly those that involve conditions that may perturb the glycans themselves would suffer from glycoform bias. Amine-labelling avoids these issues by targeting the aglycone component of glycoproteins. However, it suffers from markedly more contamination by intracellular proteins(9, 10). These contaminants reduce detection of low abundance PM proteins and obscure true PM signal by preventing discrimination between surface-localized and intracellular pools of the same protein. This is particularly problematic for heavily trafficked or recycled proteins, but is a common feature of all PM proteins since they are synthesized intracellularly(11).

The source of intracellular protein contamination in amine-labelling studies remains poorly understood, and is the focus of this study. Using amine-reactive reagents we demonstrate that biotinylation results in a small but reproducible population of intracellularly labelled cells which disproportionately contaminate downstream affinity purification of PM proteins. We further demonstrate that post-labelling depletion of Annexin V positive dead cells removes ∼99% of intracellularly labelled cells and markedly improved PM identification and peptide coverage. Moreover, using LFA-1 as an example we show that dead cell depletion selectively reduces the intracellular pool of PM proteins. Overall, we identify a major source of contamination in amine-labelling PM enrichment, establish a simple and highly effective approach to remove these contaminants, and show that this increases both the sensitivity and accuracy of surfaceomics.

### Experimental Procedures

#### Experimental Design and Statistical Rationale

This study aimed to identify the source of intracellular protein contaminants in plasma membrane (PM) enrichment proteomics from cultured cells. To investigate the origin of nonspecific intracellular contaminants, we employed a flow cytometry-based approach to evaluate multiple amine-reactive biotinylation reagents. These experiments revealed that compromised cells present during surface labeling— specifically dead or dying cells-are a primary source of intracellular contamination. To address this, we systematically compared several Annexin V-based depletion strategies to selectively remove nonviable cells prior to biotinylation. PM protein enrichment efficiency was assessed using two independent biological replicates of surface biotinylation and streptavidin-based affinity purification in Jurkat cells, followed by data-dependent acquisition (DDA) mass spectrometry on a Thermo Orbitrap Fusion Lumos. Raw data were processed with MaxQuant v2.1.3 using the human UniProt database, with a 1% false discovery rate (FDR) applied at both the peptide and protein levels(12). Figures 3 and 4 display data from a representative replicate selected for optimal streptavidin pull-down efficiency and protein yield; a second independent replicate was performed and showed consistent plasma membrane protein enrichment (Figure S3A-C). Although selected figures present single-replicate results, the reproducibility of labeling patterns across biological replicates supports the robustness and reproducibility of the findings. Statistical comparisons between experimental conditions were performed using unpaired two-tailed t-tests with Welch’s correction to account for unequal variances.

### Cell Lines and Primary Human Samples

Jurkat, HeLa, and HEK293 cells were obtained from the American Type Culture Collection (ATCC). Primary human peripheral blood mononuclear cells (PBMCs) were isolated from leukopaks (StemCell Technologies) using immunomagnetic negative selection kits (BioLegend). Jurkat and primary PBMCs were cultured in RPMI-1640 medium supplemented with 10% fetal bovine serum (FBS), 1% penicillin-streptomycin-glutamine, and 0.1% 2-mercaptoethanol. HeLa and HEK293 cells were maintained in Dulbecco’s Modified Eagle Medium (DMEM) with 10% FBS and 1% penicillin-streptomycin-glutamine. All cultures were incubated at 37°C in a humidified atmosphere containing 5% CO₂.

### T Cell Isolation and Culture

PBMCs were isolated using the EasySep Direct Human PBMC Isolation Kit (StemCell Technologies), followed by negative selection of naïve CD4⁺ T cells using the MojoSort Human CD4 Naïve T Cell Isolation Kit (BioLegend). Isolated cells were cryopreserved in RPMI supplemented with 10% DMSO and 10% FBS. For expansion, thawed CD4⁺ T cells were cultured in complete RPMI supplemented with 5 ng/mL recombinant human IL-7 and IL-15 (Fisher Scientific) for 14 days prior to biotinylation and flow cytometric analysis.

### Plasma Membrane Biotinylation

Surface protein labeling was performed using sulfo-NHS-based biotin reagents (Thermo Fisher) including sulfo-NHS-biotin, sulfo-NHS-LC-biotin, sulfo-NHS-SS-biotin, and sulfo-NHS-LCLC-biotin. Cells were washed twice with PBS supplemented with 0.5% FBS (PBS^sup^), then resuspended in PBS^sup^ at 1 × 10⁸ cells/mL. Biotin reagents (0.5 mM final concentration) were added and incubated at 4°C for 20 min. Reactions were quenched by adding 50 mM Tris-HCl (pH 8.0) and incubated for 5 min at 4°C. Cells were then washed twice in PBS and processed for flow cytometry or snap-frozen and stored at –80°C.

### Annexin V negative selection (dead cell depletion)

Following biotinylation, dead cells were removed using one of the following methods: fluorescence-activated cell sorting (FACS), Akadeum microbubble-based depletion, MojoSort Dead Cell Removal Kit (BioLegend), or the Dead Cell Removal Kit (Miltenyi Biotec)(13). FACS was performed by gating live cells on FSC-H vs. SSC-H. Manufacturer protocols were followed for all other methods. Post-depletion, viable cell counts and yields were recorded. The Miltenyi Dead Cell Removal Kit was used exclusively for all subsequent experiments after those shown in Figure 1B.

**Fig. 1.**
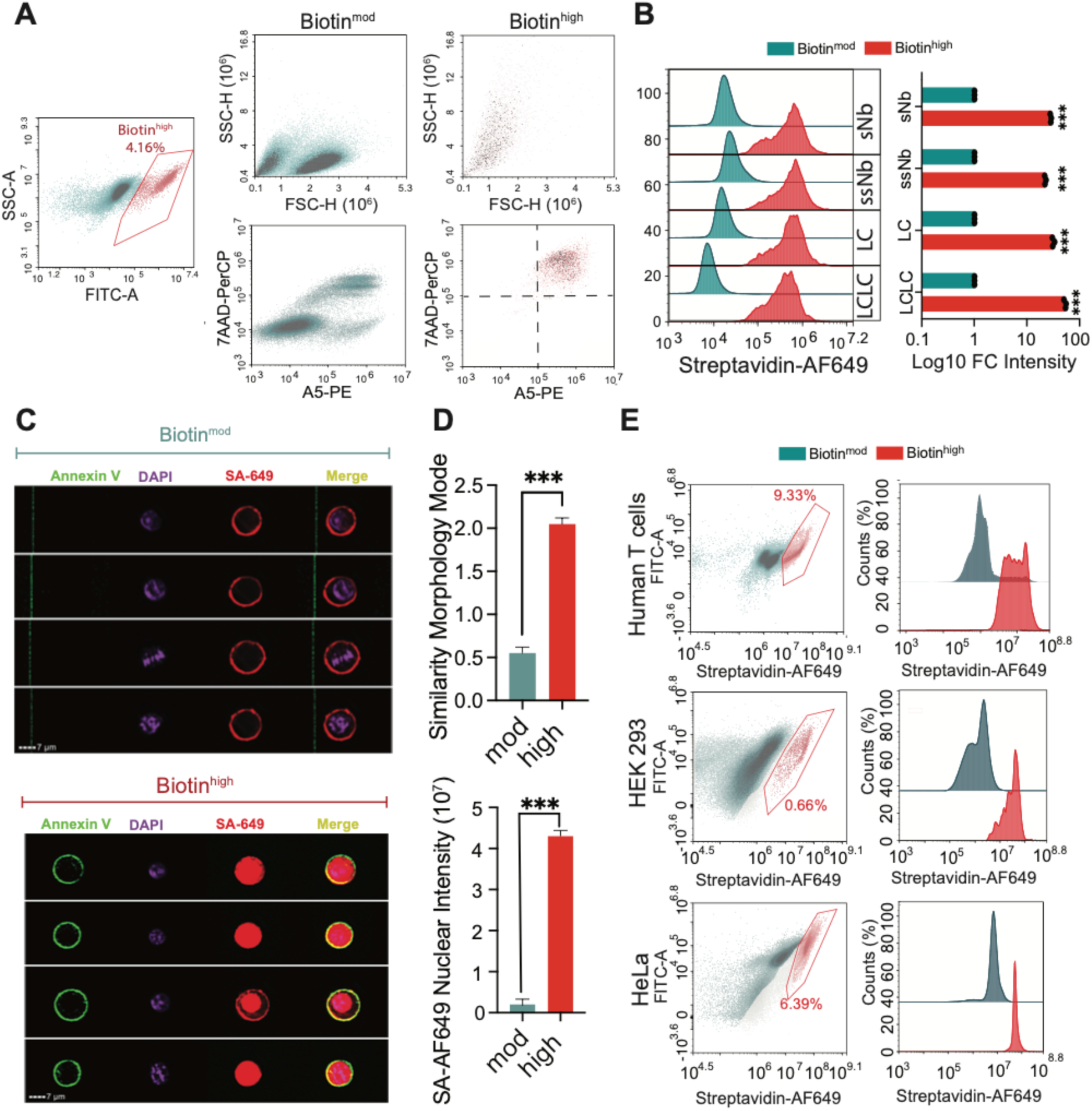
Non-Viable Cells Drive Intracellular Protein Contamination in Amine-Reactive Surface Biotinylation. A) Jurkat cells were labeled with sulfo-NHS-biotin and stained with streptavidin-FITC, 7-AAD-PerCP, and Annexin V-PE to assess biotinylation levels and cell viability by flow cytometry. Dot plots gated on biotin^high^ (right) and biotin^mid^ (middle) are shown. (B) Jurkat cells were biotinylated using one of four NHS-biotin variants: sulfo-NHS-biotin (sNb), sulfo-NHS-SS-biotin (ssNb), sulfo-NHS-LC-biotin (LC), or sulfo-NHS-LCLC-biotin (LCLC). Cells were then stained with streptavidin-AF647, and median fluorescence intensities (MFI) of biotin^mid^ and biotin^high^ populations were quantified by flow cytometry. (C) Representative images of Jurkat cells labeled with sulfo-NHS-biotin, stained with streptavidin-AF647, Annexin V, and DAPI, and acquired using the ImageStream®X Mark II imaging flow cytometer to visualize biotin localization and cell viability. (D) Cells were gated based on biotin signal intensity, and nuclear co-localization of the biotin signal (top) and nuclear streptavidin staining intensity (bottom) were quantified using IDEAS® software. (E) Human T cells, HeLa, and HEK293 cells were biotinylated with sulfo-NHS-biotin, stained with streptavidin-AF647, and analyzed by flow cytometry to examine biotin^high^ populations across cell types. ***p < 0.001, ****p-value < 0.0001 by unpaired two-tailed t-test with Welch’s correction.

### Flow Cytometry and Imaging Cytometry

Flow cytometric analysis was performed using the NovoCyte Quanteon flow cytometer. Imaging flow cytometry was carried out on the ImageStream®X Mark II system. Surface and intracellular staining was performed in triplicate using antibodies against CD11a (clone HI111) and CD18 (clone CBR LFA-1/2) from BioLegend, and Calnexin (polyclonal, Abcam). Streptavidin-DyLight 649 and Streptavidin-FITC were used for detecting biotinylated proteins (Vector Laboratories). Cell viability was assessed using Annexin V and 7-AAD (BioLegend).

### Plasma Membrane Proteomics Sample Preparation

Frozen biotinylated cell pellets were lysed in 8 M urea buffer containing 0.3 M NaCl, 50 mM sodium phosphate (pH 8.0), 0.5% IGEPAL CA-630, and 1% SDS. Lysates (1 mL per 20 million cells) were homogenized via needle shearing (19G, 21G, and 23G), followed by 30 s sonication. Lysates were centrifuged (15,000 × g, 15 min), and supernatants were incubated overnight at 4°C with streptavidin agarose resin (Thermo Fisher; 112.5 µL slurry per mL of lysate).

Beads were washed five times with lysis buffer, followed by five washes in 8 M urea wash buffer (1.22 M NaCl, 0.5% SDS, 125 mM Tris-HCl pH 8.0, 10% ethanol, 10% isopropanol). Beads were washed with 25 mM ammonium bicarbonate (ABC), resuspended in 8 M urea in 25 mM ABC, and reduced with 5 mM TCEP (37°C, 15 min), followed by alkylation with 10 mM iodoacetamide (RT, dark, 30 min). Beads were digested on-bead first with LysC (1.6 µg per 100 µL bead volume, 4 h at 37°C), then diluted to 1.3 M urea and digested with trypsin (3.2 µg per 100 µL, 14 h at 37°C). Supernatants were acidified (TFA to pH 3), desalted using C18 cartridges (pre-conditioned with ACN, 50% ACN, 0.5% acetic acid, and 0.1% TFA), and eluted with 80% ACN. Eluates were dried via vacuum centrifugation and stored at –80°C.

### Reversed-Phase Liquid Chromatography−Mass Spectrometry

Peptide mixtures were analyzed by liquid chromatography-tandem mass spectrometry (LC-MS/MS) using a Thermo Scientific™ UltiMate™ 3000 UHPLC system coupled online to a Thermo Scientific™ Orbitrap Fusion Lumos™ Tribrid™ Mass Spectrometer via a nano-electrospray ion source. Peptides were separated by reverse-phase chromatography on a 50 cm × 75 μm I.D. EasySpray Acclaim® PepMap™ RSLC C18 analytical column (Thermo Fisher Scientific) using a linear 87-minute gradient from 4% to 22% acetonitrile in 0.1% formic acid (solvent A: 0.1% formic acid in water; solvent B: 0.1% formic acid in acetonitrile) at a flow rate of 300 nL/min. Full MS scans were acquired in the Orbitrap over a mass range of 375 1500 m/z at a resolution of 60,000 (at m/z 400), followed by data-dependent MS/MS scans of the top 15 most intense precursor ions using higher collision-induced dissociation (HCD with a normalized collision energy of 30% in the linear ion trap. Dynamic exclusion was enabled with a duration of 30 seconds. The AGC target was set to 4 × 10^5^ for MS1 and 6 × 10^e^ for MS2, with maximum injection times of 50 ms and 35 ms, respectively. Each sample was analyzed in technical duplicates to ensure reproducibility.

### Data Processing

Protein quantification from LC-MS/MS data was performed using MaxQuant software (version 1.6.0.16)(12). Raw mass spectrometry files were searched against the human SwissProt database (22,355 protein entries; July 2024)(14). MS/MS spectra were filtered to retain a maximum of eight fragment peaks per 100 Da interval. The precursor mass tolerance was set to 20 ppm for the initial search and 4.5 ppm for the main search. False discovery rates (FDR) for both peptide spectral matches and protein identifications were controlled at 1%, using the razor peptide approach. Trypsin was specified as the digestion enzyme, allowing up to two missed cleavages, with no allowance for nonspecific cleavage. Carbamidomethylation of cysteine residues was defined as a fixed modification, while methionine oxidation and N-terminal acetylation were set as variable modifications, allowing up to two variable modifications per peptide. Protein intensities were calculated as the integrated peak area across the chromatographic elution profile. For isotopic clusters, intensities from all isotopic peaks were summed. Only peptides classified as unique plus razor were used for quantification to ensure protein-level accuracy.

### Western Blotting

For SDS-PAGE, bead-bound proteins were eluted by boiling (95°C, 20 min) in 1× LDS sample buffer (Fisher Scientific). Supernatants were collected following centrifugation (1600 rpm, 5 min). Input and flow-through samples were quantified by UV absorbance and normalized to 20 µg per lane. Proteins were resolved on 4–12% Bis-Tris gels (Thermo Fisher) and transferred to membranes for immunoblotting. Antibodies were used against integrin β2 (D4N5Z), integrin αL (E5S9K), CD45 (D9M8I), CD7 (E4G1Q), Histone H3 (polyclonal), and β-actin (13E5) (Cell Signaling Technology), as well as PCCA (polyclonal, Abcam). Imaging was performed on an iBright FL1500 system (Invitrogen).

### Streptavidin Bead Flow Cytometry

Beads were washed twice with PBS, passed through a 70 µm cell strainer, centrifuged (400 × g, 10 min), and plated in 96-well round-bottom plates. Staining was performed using the same protocol as for cell-based flow cytometry. Antibodies against β-actin (SP124) and histone H3 (EPR16987) were obtained from Abcam.

### On-Bead Endoglycosidase Treatment

To assess glycoprotein status, streptavidin-bound proteins were treated with Endo H or PNGase F (New England Biolabs) following three PBS washes. Reactions were performed according to the manufacturer’s protocols. Beads were then washed with lysis buffer and processed for western blotting

## Results

### Amine-labelling produces dead cells with high intracellular protein labels

For poorly understood reasons, cell surface labelling and enrichment using lysine (amine) reactive biotinylation reagents results in contamination with intracellular proteins(15–19). Using flow cytometry to optimize cell surface biotinylation conditions of Jurkat T cells, we observed a bimodal distribution of streptavidin staining intensity (Fig. 1A). While ∼95% of cells were moderately labelled (biotin^mod^ cells), a small but distinct and reproducible population showed mean fluorescence intensity (MFI) values that were ∼20 to 100-fold higher (biotin^high^ cells). Scatter profiles indicated that biotin^high^ cells had decreased forward scatter and increased side scatter, features consistent with non-viable Jurkat T cells. To directly examine cell viability, we performed 7-aminoactinomycin D (7-AAD) and Annexin V staining. 7-AAD is a membrane impermeable DNA-binding dye, which positively stains cells that have lost membrane integrity. Biotin^high^ cells were positive for both markers, confirming compromised membrane integrity, and identifying them as dead or dying (Fig. 1A). To determine whether this labeling pattern is a general feature of amine-labelling reagents or unique to sulfo-NHS-biotin, we also tested the amine-reactive reagents sulfo-NHS-SS-biotin, sulfo-NHS-LC-biotin, and sulfo-NHS-LC-LC-biotin. All reagents tested resulted in the appearance of a biotin^high^ cell population at similar frequencies (Fig. S1A). Fluorescence intensities of the biotin^high^ population remained approximately 20- to 100-fold higher than the biotin^mod^ population across all reagents (Fig. 1B).

Given the much higher abundance of intracellular proteins compared to PM proteins, the markedly higher labelling in these dead cells is consistent with intracellular biotinylation. To confirm and visualize intracellular labelling, we compared staining patterns in biotin^mod^ and biotin^high^ cells by imaging flow cytometry. Jurkat cells labeled with sulfo-NHS-biotin and stained with streptavidin-DyLight649, Annexin V-FITC, and the nuclear marker DAPI were imaged and analyzed. In biotin^mod^ cells, streptavidin signal localized predominantly to the PM. In contrast, biotin^high^ cells exhibited strong intracellular staining, with pronounced nuclear localization in many cells (Fig. 1C and S1B). Co-localization analysis between streptavidin-DyLight649 and DAPI signals revealed a five-fold increase in similarity score in biotin^high^ cells, further supporting nuclear enrichment. Quantitative measurement using a DAPI-defined nuclear mask also showed a nearly five-fold increase in nuclear streptavidin signal in biotin^high^ versus biotin^mod^ cells (Fig. 1D).

We next assessed whether the intracellular biotinylation occurs in other cell types, including primary lymphocytes cells and adherent cells. Primary human CD4⁺ T cells isolated from leukopaks exhibited an even higher frequency of biotin^high^ cells (∼9%), likely due to reduced viability from sample transport and handling (Fig.1E). Similarly, adherent HEK293 and HeLa cells labeled with sulfo-NHS-biotin also showed biotin^high^ populations, though the frequency of this population varied (Fig. 1E). These findings suggest that intracellular labeling of non-viable cells is a general phenomenon of amine-reactive biotinylation protocols across diverse cell types. Given the ∼20-100 fold higher labelling observed in these events, our data also suggest that a small population of non-viable cells can contribute disproportionately to contamination. For example, intracellularly labelled cells accounting for just 2% of the population, but displaying 50-fold higher labelling, would be expected to contribute as much protein as the remaining 98% of live, PM-labelled cells. These findings are consistent with previous reports highlighting significant intracellular and nuclear contamination in amine-labelling surface proteomics workflows (20, 21).

### Biotin^high^ cells can be efficiently depleted post-biotinylation

We next examined whether altering labelling conditions or cell culture conditions could reduce the biotin^high^ population. Titrating reagent concentrations revealed that the biotin^high^ population was detectable even when biotinylating at 0.0625 mM for all biotin reagents tested (Fig. S2A). Similarly, reducing the biotinylation reaction time resulted in a modest decrease in fluorescence intensity but did not significantly alter the percentage of biotin^high^ cells (Fig. S2B). Finally, we examined the impact of improving cell viability by reducing cell culture density, on the frequency of biotin^high^ cells. ATCC recommends growing Jurkat cells up to 3×10^6^ cells per ml. We routinely culture Jurkat cells up to 2×10^6^ cells per ml. As expected, increasing cell density further led to more cell death and increased proportions of biotin^high^ cells. Reducing cell culturing density moderately reduced the proportion of biotin^high^ cells, but failed to eliminate them entirely (Fig S2B).

We thus turned our attention to removal of these cells post-labelling. Since biotin^high^ cells are Annexin V positive (Fig. 1A), we hypothesized that Annexin V based negative selection could effectively deplete dead cells and reduce intracellular labeling. Commercially, several Annexin V based methods are available for dead cell removal. In addition, as shown previously (Fig. 1A), biotin^high^ cells exhibit distinct FSC/SSC profiles, enabling separation by fluorescence-activated cell sorting (FACS)(23). FACS sorting based on FSC/SSC alone efficiently reduced biotin^high^ cells from 4.5% to 0.23%. Although additional sorting based on viability markers would likely improve this further, FACS sorting was slow, required specialized equipment, and yielded low recovery (∼25%) (Fig. 2B).

**Fig. 2.**
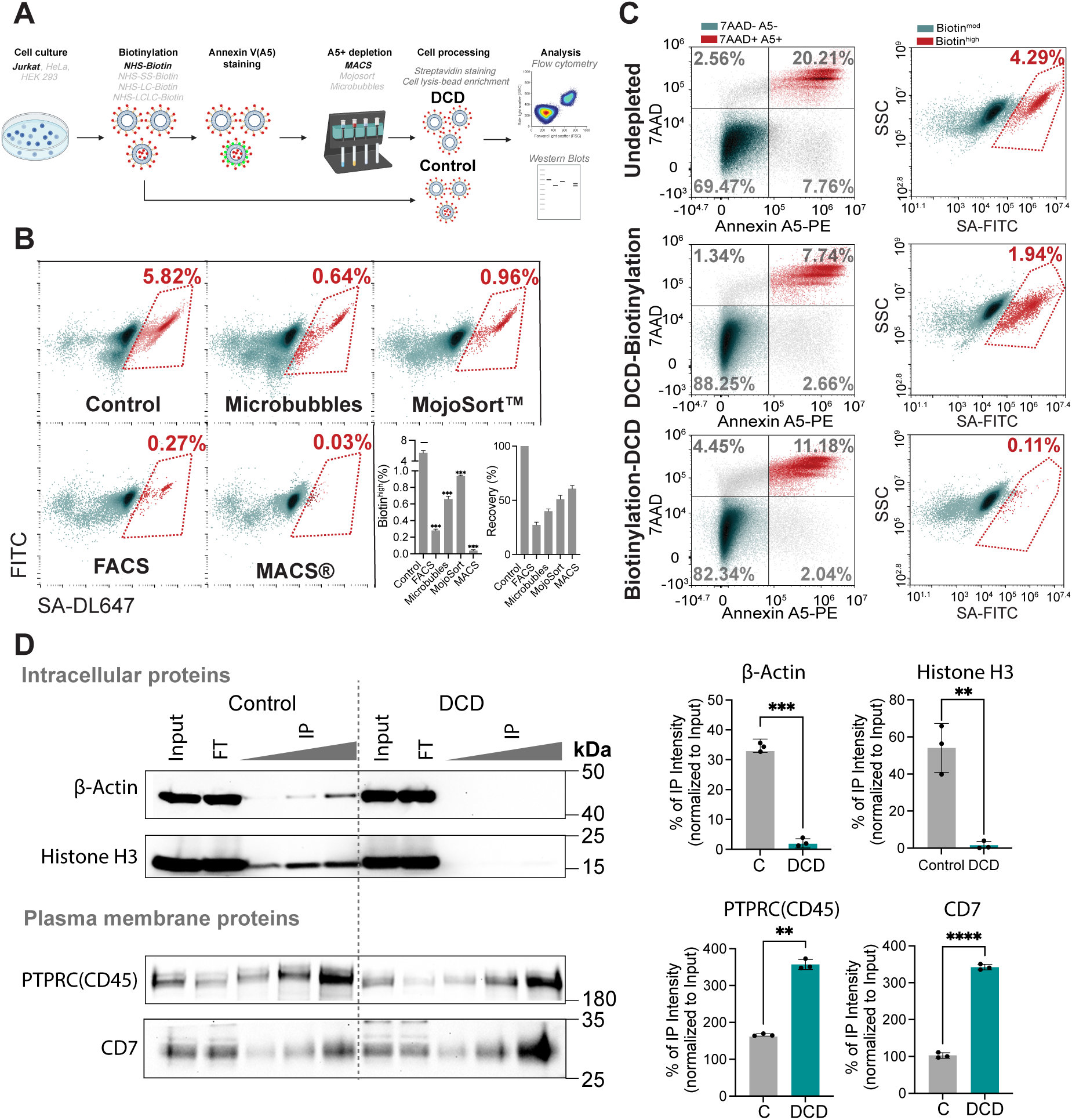
Optimization of Dead Cell Removal Strategies to Minimize Intracellular Protein Contamination Following Amine-Reactive Biotinylation. (A) Schematic illustration of the experimental workflow, depicting NHS-biotin surface labeling followed by dead cell depletion. (B) Annexin V–positive cells remaining after NHS-biotin surface labeling were depleted using various methods as indicated. The efficiency of depletion and cell recovery rates were quantified by flow cytometry. (C) Jurkat cells containing Annexin V–positive cells were either left undepleted (top panel), depleted prior to surface labeling (middle panel), or depleted after surface labeling (bottom panel). Cells were stained with streptavidin to assess the efficiency of removing the Biotin^High^ population by flow cytometry. (D) Surface-labeled cells, with or without Annexin V–positive cell removal, were lysed, and biotinylated proteins were enriched using streptavidin-coated agarose beads at 4°C overnight. Post-enrichment, proteins were eluted and loaded onto SDS-PAGE gels at varying volumes (10 μL, 20 μL, and 40 μL), alongside input and flow-through (FT) fractions. Western blotting was performed using primary antibodies against Actin, Histone H3, PTPRC (CD45), and CD7, followed by HRP-conjugated secondary antibodies. Band intensities were quantified using ImageJ, and immunoprecipitated (IP) signals were normalized to input to calculate the percentage of enrichment. NS, not significant; *p < 0.05; **p < 0.01; ***p < 0.001 by unpaired two-tailed t-test with Welch’s correction.

We therefore turned our attention to three commercially available Annexin V– based kits: Miltenyi magnetic beads (MACS), Akadeum microbubbles, and the BioLegend Dead Cell Removal Kit. These approaches have the advantage of processing large numbers of cells quickly, increasing potential suitability for downstream proteomics. Among them, MACS achieved the highest depletion efficiency, reducing biotin^high^ cells from 4.5% to 0.03%, with ∼60% recovery. The BioLegend kit reducing dead cells to 0.95%, while the microbubble method showed better depletion but also more cell loss (∼40% recovery) (Fig. 2B). Given the highly effective depletion of biotin^high^ cells, short processing times, and relatively high cell recovery, we proceeded with the MACS approach as our standard dead cell depletion method.

We then compared pre-versus post-biotinylation dead cell depletion. While depleting dead cells prior to sulfo-NHS-biotin labeling modestly reduced the biotin^high^ population, it was notably less effective than dead cell depletion post-labelling (Fig. 2C). This suggests that the labeling itself or the general processing involved continually induce cell death. Indeed, a similar number of 7-AAD⁺Annexin V^+^ cells persisted in both conditions. We also examined the effectiveness of dead cell depletion in primary human T cells. Growing primary human T cells in the presence of the pro-survival cytokines IL-7 and IL-15 for two weeks reduced the number of biotin^high^ cells to ∼2%. However, similar to the Jurkat cells, dead cell depletion using the MACS method, reduced this number further to 0.06%, essentially eliminating biotin^high^ events (Fig. S2C). Overall, these data demonstrate that sulfo-NHS-biotin labeling bypasses the PM in a subset of cells and that post-labeling dead cell depletion can efficiently remove these intracellularly labelled biotin^high^ events.

### Dead cell depletion reduces intracellular contaminants and improves surfaceomics

We expected the removal of intracellularly biotinylated cells to deplete intracellular contaminants and improve downstream enrichment of cell surface proteins. To first estimate the extent of this improvement on surface proteome quality, we compared streptavidin bead capture (IP) for key intracellular and PM proteins from NHS-biotin-labeled samples, with or without dead cell depletion. Post-labeling dead cell depletion dramatically reduced the detection of histone H3 and β-actin by western blotting of captured proteins, while enhancing the enrichment of the known Jurkat plasma membrane proteins CD7, CD45 and integrin β2 (Fig. 2D and S2D).

Next, we evaluated the effect of dead cell depletion on cell surface proteomic profiling by mass spectrometry. Dead cell depletion resulted in fewer total proteins (1,599 versus 3,033) and peptides (12,687 versus 24,157) than the control sample (Fig. 3A-B, Table S1). Despite this however, it markedly enhanced enrichment of PM proteins. Using SURFY annotations (Table S2) (7), 281 PM proteins were identified with dead cell depletion, compared to 152 in the control sample, representing an 85% increase. As a proportion of total ID’s, bona fide PM ID’s increased from 5.1% to 17.6% with dead cell depletion (Fig. 3C). In addition, the PM proteins identified had dramatically higher peptide coverage and were present at much higher intensities (Fig. 3D-E). Concurrently, the peptide coverage and intensities of non-PM proteins were markedly reduced with dead cell depletion consistent with reduced intracellular contamination (Fig. 3D-E). A second independent surfaceomics replicate was performed and showed consistent plasma membrane protein enrichment (Figure S3A-C).

**Fig. 3.**
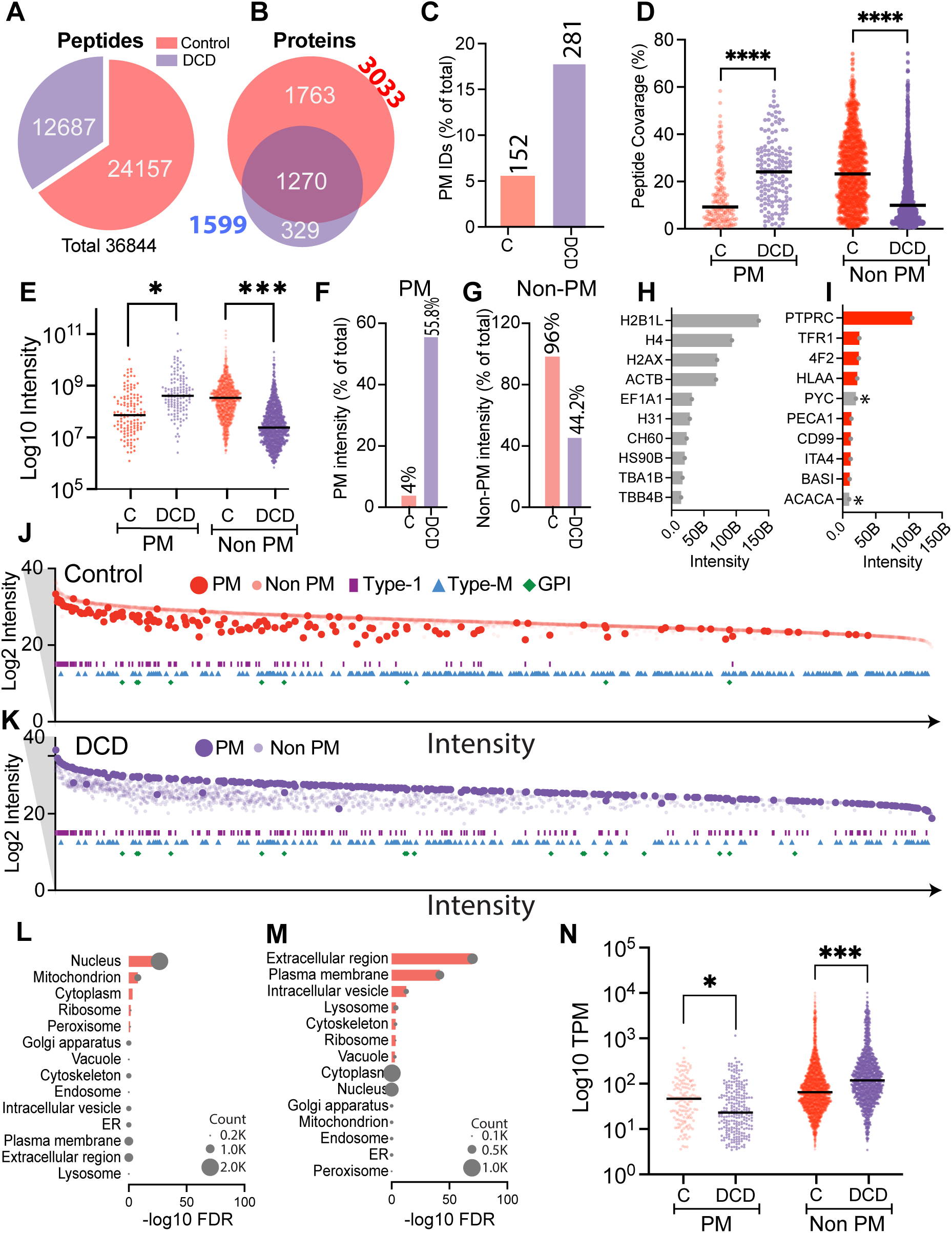
Annexin V–Based Dead Cell Depletion Enhances Specificity and Sensitivity of Plasma Membrane Proteomics by Mass Spectrometry. Peptide (A) and protein (B) overlaps between control and DCD runs are shown. (C) Quantification of plasma membrane (PM) proteins in control and DCD samples was performed based on SURFY annotations. Peptide coverage percentages for PM and non-PM (D) and iBAQ intensities (E) are compared between control and DCD samples in (D). iBAQ intensities. The percentage intensity of PM proteins (F) and Non-PM proteins (G) was calculated relative to the total protein intensity within each sample. Comparisons were made between control and DCD groups to assess distribution differences. Bar plots illustrate the top 10 highest-abundance proteins based on raw intensity in control (H) and DCD (I) samples, with red bars indicating SURFY-annotated PM proteins and asterisks denoting endogenous biotinylated proteins. (J, K) Scatter plots depict proteins identified in control (J) and DCD (K) datasets, where PM proteins are shown as large, dark circles (red in control, blue in DCD) and non-PM proteins as smaller, lighter circles. Protein features are annotated as purple rectangles for type I membrane proteins, cyan triangles for multi-pass proteins, and green diamonds for GPI-anchored proteins. The x-axis represents summed intensities across control and DCD datasets, and the y-axis indicates intensities in either control or DCD samples. (L, M) Subcellular enrichment analyses were performed using SubcellulaRVis with the Jurkat genome as background, with red bars indicating significantly enriched compartments and bubble sizes reflecting the number of proteins identified in control (L) and DCD (M) samples. (N) Comparison of Human Protein Atlas–derived expression levels is shown for PM and non-PM proteins identified in control and DCD samples. NS, not significant; *p < 0.05; **p < 0.01; ***p < 0.001 by unpaired two-tailed t-test with Welch’s correction.

As an aggregate measure of PM enrichment, we examined the cumulative proportional intensities of all PM and non-PM proteins. The aggregate proportional intensity of all PM proteins increased from 4% in the control sample to 55.8% with dead cell depletion, indicating a ∼14-fold enrichment in PM proteins (Fig. 3F). By contrast, total intracellular protein intensity decreased from 96% to 44.2% (Fig. 3G).

We next examined the top 10 most abundant proteins in each condition. Without dead cell depletion, the top-ranking proteins were predominantly high-abundance intracellular contaminants such as nuclear histones and cytoplasmic actin (Fig. 3H). In contrast, the most abundant proteins with dead cell depletion were PM-localized, with the exception of two endogenously biotinylated mitochondrial proteins (ACACA and PYC) (Fig. 3I) (24, 25). These findings suggest that without dead cell depletion, proteomic profiles are biased toward abundant intracellular contaminants, obscuring true surface protein enrichment.

To visualize broader trends in proteome composition, we mapped all quantified proteins by summed intensity and annotated structural features. In dead cell depleted samples, PM proteins formed a distinct and concentrated cluster, in stark contrast to the scattered distribution observed in control samples (Fig. 3J-K). Additionally, dead cell depletion produced a greater enrichment of Type I transmembrane proteins (purple rectangles) and GPI-anchored proteins (green diamonds) (Fig. 3J-K, Table S2). Multi-pass transmembrane proteins (blue triangles) were comparably represented in both conditions. These results indicate that dead cell depletion preferentially enriches proteins with canonical cell surface features (14). The lower enrichment of multipass membrane proteins in our dataset may be attributable to the high hydrophobicity of their transmembrane peptides, which can reduce solubilization efficiency and peptide recovery during sample preparation and LC-MS/MS analysis (26, 27).

Gene Ontology (GO) enrichment analysis further supported these findings. The control data set was significantly enriched for nuclear, cytoplasmic, and mitochondrial proteins, indicative of pervasive intracellular contamination (Fig. 3L) (14, 28). Conversely, dead cell depletion demonstrated strong enrichment for proteins localized to the plasma membrane, extracellular space, and endosomal compartments, with high statistical significance (FDR < 0.05) (Fig. 3M). These shifts confirm the enhanced specificity of PM proteomes derived by dead cell depletion (14, 28).

To investigate whether transcript abundance influences protein detection in our datasets, we analyzed gene expression profiles of Jurkat E6-1 cells using publicly available RNA-seq data from the Human Protein Atlas (Table S1) (29). PM proteins uniquely identified in control samples exhibited significantly higher transcript levels compared to those exclusively detected in Annexin V-based dead cell–depleted samples. This suggests that in control samples, highly expressed PM proteins dominate the profile, potentially due to signal dilution from substantial contamination by abundant intracellular proteins. In contrast, the reduction of intracellular proteins by dead cell depletion facilitated the detection of lower-abundance PM proteins (Fig. 3N). Among the identified PM proteins, those found only with dead cell depletion had significantly lower transcript levels compared to PM proteins detected in control samples or in both conditions (Fig. S3D). Collectively, these data demonstrate that Annexin V–based dead cell depletion significantly improves both the specificity and sensitivity of plasma membrane proteomic profiling by reducing intracellular contamination and enhancing the detection of lower-abundance surface proteins.

### Quantitative scoring reveals specific enrichment of surface-localized proteins with dead cell depletion

To better characterize the specificity of PM protein enrichment with dead cell depletion, we calculated a quantitative enrichment score based on the ratio of protein intensities between dead cell depleted and control samples. This metric allowed us to assess the degree of enrichment for each protein across conditions. Among 129 SURFY annotated PM proteins identified in both samples, the mean enrichment score was 5.7. Notably, 120 of these proteins had an enrichment score greater than 1, indicating increased abundance with dead cell depletion. The remaining nine proteins with scores <1 were either intracellular proteins or proteins with ambiguous localization. For instance, EBP, PO210, RPN1, STT3B, and ATP13A1 are predominantly intracellular, while ATP1A1, ULBP2, and TM9SF3 are annotated as both PM and intracellular in various databases (Fig. 4 A, Table 1S) (30–33). In contrast, proteins annotated as non-PM had a mean enrichment score of 0.06. Of these, 1,045 had scores <1, and only 96 exceeded 1 further supporting selective depletion of intracellular proteins. Interestingly, two proteins with the highest enrichment scores, BPL1 (score = 66.6) and ACACA (score = 54.6) are known to be endogenously biotinylated (BPL1 catalyzes the biotinylation of carboxylases and histones, while ACACA is a biotin-containing enzyme), likely accounting for their elevated scores (34, 35). Excluding such exceptions, approximately 20 high-scoring proteins not annotated by SURFY were found to be PM proteins in other databases (14, 36), including well established PM proteins such as transferrin receptor (TFR1) and components of the T cell receptor complex (TRAC, TRBC1 and TVBL3) (Fig. 4A). These findings suggest that despite annotation ambiguities, dead cell depletion effectively enriches bona fide surface proteins.

**Fig. 4.**
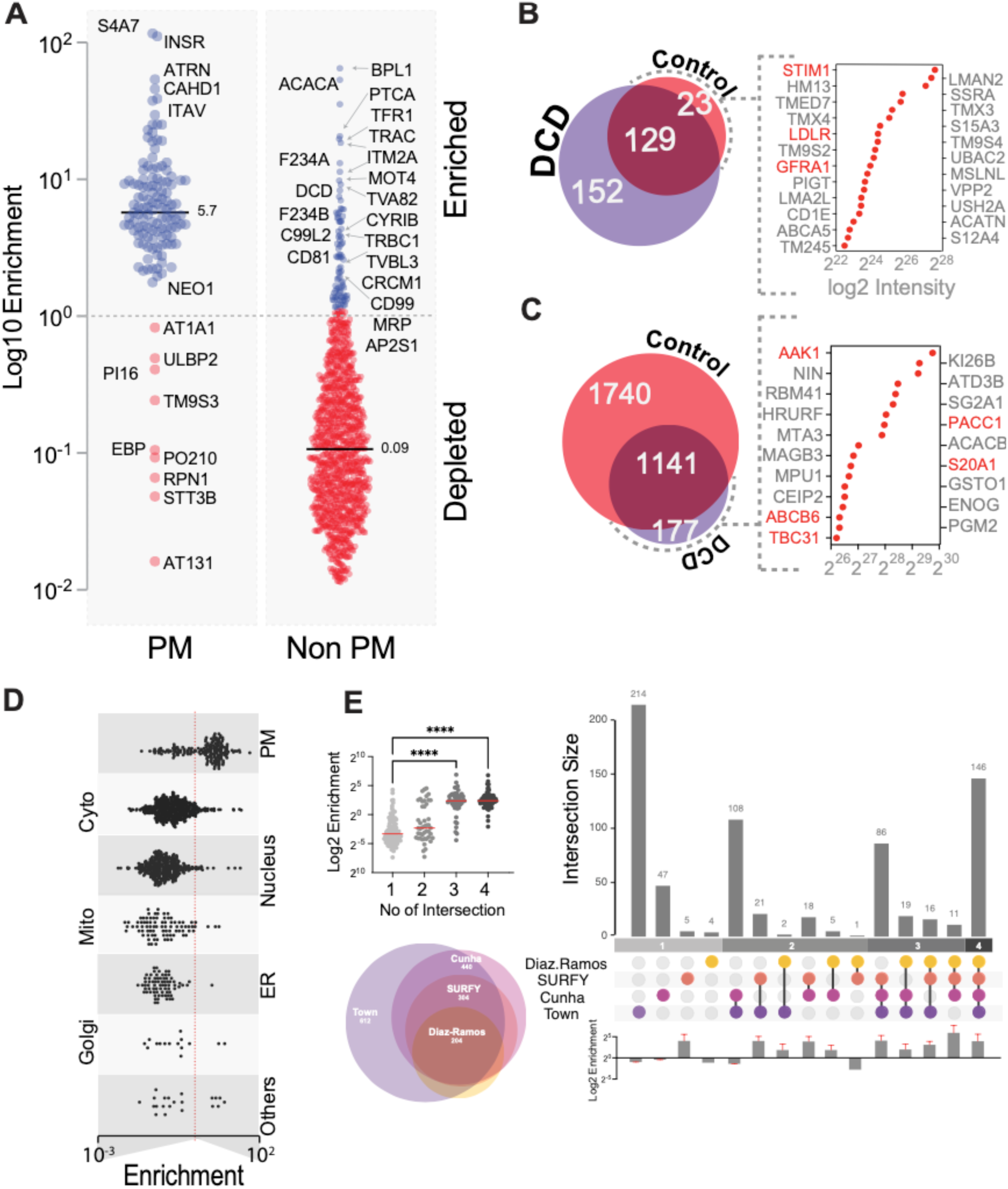
Quantitative Enrichment Analysis Demonstrates Improved Specificity for Plasma Membrane Proteins Following Annexin V Dead Cell Depletion. (A) Scatter plot displays enrichment scores calculated as the ratio of raw intensities in DCD versus control samples, with proteins annotated according to SURFY, where blue circles represent proteins with enrichment scores greater than 1 and red circles represent scores less than 1. (B) The overlap of SURFY-annotated PM and non-PM proteins between control and DCD samples is shown; the top right scatter plot displays raw intensities of PM proteins uniquely detected in control samples, while the bottom right scatter plot shows raw intensities of non-PM proteins uniquely detected in DCD samples. (C) Enrichment scores for proteins categorized by subcellular localization are compared based on UniProt annotations. (D) Multiple plots integrate PM annotation confidence across databases: the top left histogram depicts the number of proteins annotated in one or more databases alongside their enrichment scores; the bottom left Venn diagram shows overlaps of proteins identified in both control and DCD datasets and their presence in different PM annotation resources; the Upset plot visualizes intersections of proteins identified in both datasets and annotated as PM in varying combinations of databases; and the bottom histogram presents enrichment scores for each combination of database overlaps.

We next examined proteins that were only identified in the control or dead cell depletion samples. The 152 unique proteins with SURFY PM annotation in the dead cell depleted sample were also consistently annotated as PM proteins in other databases (14, 36). By contrast, of the 23 unique proteins with SURFY PM annotation in the control sample (Fig 4B, Table S3), the majority were consistent with non-PM localization (7, 14). Conversely, 177 unique proteins annotated as non-PM by SURFY were identified in the dead cell depleted sample, of which ∼30 were nevertheless localized to or associated with the PM (7, 14). These findings further reinforce the specificity of the dead cell depletion strategy (Fig. 4C, Table S3).

Given the variability in PM annotations across databases, we next evaluated the subcellular localization of proteins using UniProt (14). We plotted enrichment scores by UniProt annotations and found that PM-localized proteins had the highest mean enrichment score (6.7), whereas proteins localized to the cytosol, nucleus, mitochondria, ER, or Golgi had mean scores <1 (Fig. 4D, Table S4). This result is consistent with selective PM enrichment in dead cell depleted samples and reduced contamination from intracellular compartments. To assess whether enrichment scores correlated with annotation confidence, we integrated four PM databases (Diaz-Ramos, Bausch-Fluck, Town and Cunha) and categorized proteins based on the number of databases supporting their PM localization (7, 37–39). We identified 454 PM proteins present in our dataset that were annotated in at least one of the databases. Proteins annotated in only one database (n = 113) had a mean enrichment score of 1.6. Those annotated in two databases (n = 79) had a mean score of 5.6; three databases (n = 119) yielded a score of 8.4; and proteins supported by all four databases (n = 79) had a mean enrichment score of 8.3 (Fig. 4E, Table S4). This trend suggests a positive correlation between enrichment score and the confidence level of PM localization. Taken together, these results indicate that dead cell depletion not only improves sensitivity for detecting PM proteins, but also selectively enriches high-confidence surface-localized proteins.

### Dead cell depletion specifically removes the intracellular pool of PM proteins

PM proteins reside in both intracellular and cell-surface pools, and distribution between these compartments can be dynamically regulated (40). For example, glucose transporters are stored intracellularly in the absence of insulin, but are quickly trafficked to the cell surface in response to insulin signaling (41). Based on our findings that sulfo-NHS-biotin labels intracellular proteins in biotin^high^ dead cells, we hypothesized that routine labelling will capture both cell-surface and intracellular pools of PM proteins and that dead cell depletion will selectively remove the intracellular contribution to more faithfully reflect the cell surface proteome. To test this hypothesis, we examined Lymphocyte Function-Associated Antigen 1 (LFA-1). LFA-1 is a heterodimer composed of integrin αL (CD11a) and integrin β2 (CD18) (42). It is a critical cell adhesion molecule expressed on leukocytes which has also been shown to have an intracellular pool in T cells (43). We first tested whether cell surface and intracellular pools could be detected in Jurkat T cells by imaging flow cytometry. Indeed, both CD11a and CD18 demonstrated extensive intracellular staining and showed at least partial co-localization with the ER marker Calnexin (Fig. 5A-B).

**Fig. 5.**
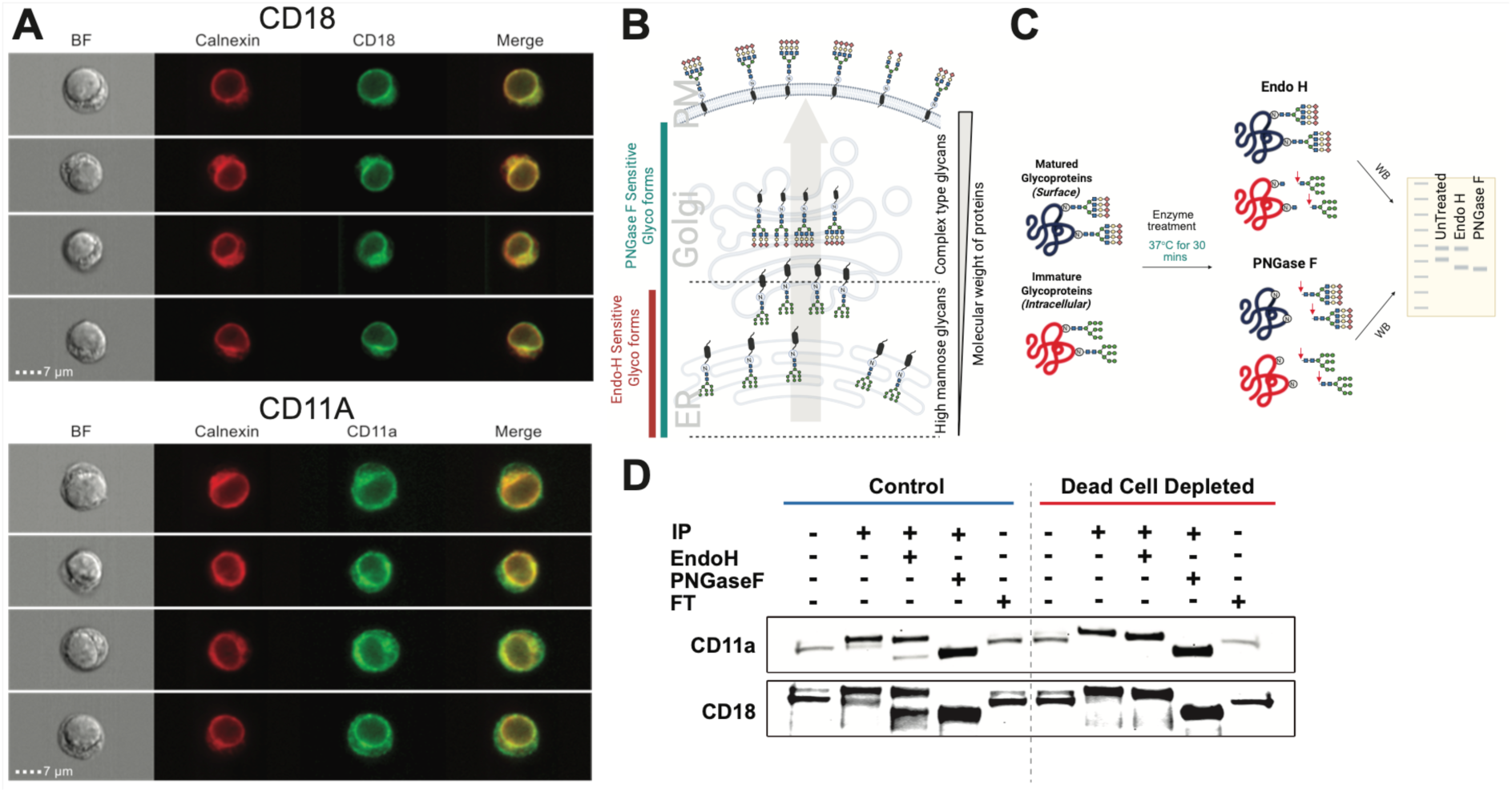
Dead Cell Depletion Selectively Removes Intracellular Pools of Plasma Membrane Proteins to Enhance Surfaceome Specificity. (A) Representative images of Jurkat cells showing intracellular localization of CD18 (ITGB2) and CD11A (ITGAL) in relation to the ER marker calnexin. Cells were fixed, permeabilized, and stained with CD18-FITC or CD11A-FITC, and calnexin, then analyzed using the ImageStream®X Mark II imaging flow cytometer to visualize subcellular distribution. (B) Schematic illustration of N-glycan maturation along the secretory pathway, highlighting the presence of Endo H–sensitive (ER-resident) and Endo H–resistant (post-Golgi) glycoforms. (C) Overview of the experimental strategy used to distinguish intracellular versus plasma membrane pools of biotin-enriched proteins. (D) Dead cell depletion reduces intracellular contamination in the biotin-enriched fraction. Biotin-labeled proteins from control and DCD samples were enriched using streptavidin agarose beads and treated with either Endo H or PNGase F overnight at 37 °C. Post-treatment, proteins were eluted and analyzed by SDS-PAGE followed by Western blotting using primary antibodies against CD18 and CD11A, and HRP-conjugated secondary antibodies.

To distinguish cell surface and intracellular pools, we leveraged selective deglycosylation by Endo H, an endoglycosidase which specifically cleaves immature N-glycans (high mannose and hybrid N-glycans) but not Golgi remodeled N-glycans (complex N-glycans) (Fig. 5C). As a control, we also used PNGase F, which cleaves essentially all N-glycans. Both CD11a and CD18 demonstrated two distinct bands by western blotting (Fig. 5D) (44). In control whole cell lysate (input), the lower molecular weight (MW) species was more abundant. However, following cell surface enrichment, the higher MW band became more abundant. This is consistent with the lower molecular weight species representing the large intracellular ER pool while the higher molecular weight band representing the cell surface glycoform that is enriched with cell surface biotinylation. Indeed, while PNGase F collapsed both bands to a lower molecular weight deglycosylated form (Fig. 5D, lane 4), Endo H only deglycosylated the lower molecular weight species (Fig 5D, lane 3), confirming that the lower MW band represents an immature glycoform. Critically, comparison of the streptavidin enriched IP fractions demonstrated that dead cell depletion results in selective loss of the Endo H sensitive, lower molecular weight bands of both CD11a and CD18 (Fig. 5D, lanes 7-9) (45, 46). These data provide proof of concept that intracellular labelling in biotin^high^ cells leads to contamination of the subsequently enriched surfaceome with intracellular sources of PM proteins, and that dead cell depletion can remove these contaminants to more accurately reflect the surfaceome.

## Discussion

Amine-reactive chemical labeling approaches, such as sulfo-NHS-biotin, are widely used in cell surface proteomics due to their ability to covalently modify accessible primary amines on extracellular lysines (15, 16, 19, 47). However, our data demonstrate that these reagents also biotinylate intracellular proteins in non-viable cells, introducing a major source of contamination. Despite comprising a small fraction of the total cell population, non-viable biotin^high^ cells exhibit 20- to 100-fold higher biotin signal than viable cells due to the substantially greater abundance of intracellular proteins (48) (Fig.1). This disproportionate labeling can dominate downstream surface-enriched proteomes, obscuring the detection of bona fide plasma membrane (PM) proteins (Fig. 2-3). While optimization of cell culture conditions and pre-labeling depletion of dead cells can modestly reduce biotin^high^ contamination, our findings highlight the superior efficacy of post-labeling dead cell depletion (Fig. 2). Removing Annexin V+ cells after sulfo-NHS-biotin labeling essentially eliminates the biotin^high^ population and significantly reduces intracellular protein contamination, as confirmed by decreased abundance of canonical intracellular markers such as histones and actin and improves the detection of transmembrane and GPI-anchored PM proteins (Fig. 2-3). This strategy increases not only the number of uniquely identified PM proteins but also enhances their peptide coverage and signal intensity, thereby improving both the specificity and quality of surfaceome analyses (Fig. 3).

Importantly, we show that intracellular labeling of dead cells includes not just cytoplasmic and nuclear proteins, but also intracellular pools of PM proteins (Fig. 5). This underscores a key limitation of surfaceome studies that rely solely on compartment-based annotation for filtering (49, 50). While annotation-based filtering can remove strictly intracellular proteins, it cannot distinguish between intracellular and surface-resident pools of proteins with dual localization, such as trafficking PM proteins (GLUT4) or proteins enroute through the secretory pathway (TFR1) (41, 51). Our quantitative enrichment scoring further supports the ability of Annexin V based depletion to enrich for high-confidence PM proteins, particularly those supported by multiple surfaceome databases (7, 37–39). Conversely, many proteins detected exclusively in control samples appear to be misannotated or derive from abundant intracellular compartments, highlighting the potential for dead cell depletion to refine surface protein annotations and reduce false positives in surfaceome datasets (Fig. 4) (7, 37–39).

Although our study specifically investigates amine-reactive labeling using sulfo-NHS-biotin, the findings may have broader implications. Many labeling chemistries, whether targeting lysines, thiols, or glycans rely on intact membrane integrity to ensure surface specificity (52). Intracellular contamination is, therefore, a general concern in chemical surface labeling and should be examined. The extent of contamination may depend on the distribution of reactive functional groups, which are typically more abundant intracellularly due to higher total protein content. For instance, lysine residues are proportionally more prevalent in intracellular proteins, which likely explains the strong intracellular signal in biotin^high^ cells (53). In contrast, glycan-based CSC strategies may show less intracellular labeling, as complex glycans are largely absent from the cytoplasm and nucleus (54). However, even glycan-reactive labels are not immune to contamination from intracellular pools of PM proteins. Indeed, glycan-based datasets frequently include ER- and Golgi-resident glycoproteins such as OST complex members and glycosyltransferases, suggesting that non-viable cells may contribute artifacts even in these workflows (55). We also demonstrate that the frequency of biotin^high^ cells can vary depending on cell type, culture/labelling conditions, and possibly other experimental details. Our simple method involving flow cytometry and/or imaging flow cytometry can be used to evaluate the extent of dead cell labelling in other experimental systems and workflows. As the percentage of biotin^high^ cells and their MFI increases, so does the expected value of dead cell depletion.

Our findings highlight an important consideration for quantitative surfaceomics, particularly in studies aiming to compare cell surface expression across conditions. Without dead cell depletion, dynamic changes in surface abundance may be masked by variable levels of intracellular contamination, especially for proteins with both surface and intracellular pools. This can result in misleading interpretations of differential expression or trafficking dynamics. Dead cell depletion is thus expected to enhance the quantitative accuracy of surface expression changes by decreasing the contribution of intracellular pools of PM proteins.

In summary, we identify post-labeling dead cell depletion as a critical step for improving the quality of amine-based surface proteomics. By effectively removing biotin^high^ non-viable cells, Annexin V-based negative selection enhances the detection of true PM proteins, reduces intracellular contamination, and increases the reliability of both qualitative and quantitative analyses. As surfaceome studies continue to expand in scale and application from immunophenotyping to biomarker and therapeutic target discovery, standardizing approaches to mitigate dead cell-driven artifacts will be essential (22, 56). Furthermore, extending these findings to other labeling chemistries, will be an important avenue for future investigation aimed at further improving cell-surface-specific proteomic profiling.

## Acknowledgments

This work was supported by the National Institute of Allergy and Infectious Diseases (DP2AI164301, H.M.). The funders had no role in the study design, data collection and analysis, decision to publish or preparation of the manuscript. We extend special thanks to Dr. Clinton Yu and Dr. Lan Huang at the High-End Mass Spectrometry Facility, UC-Irvine, for their expert assistance with mass spectrometry data acquisition and analysis. We are also grateful to Dr. Michael Demetriou and members of the Mkhikian lab for thoughtful feedback on the manuscript.

## Author Contributions

**Christofer Daniel Sánchez:** Conceptualization, Methodology, Validation, Formal analysis, Investigation, Writing - Original Draft, Visualization, Writing - Review & Editing. **Aswath Balakrishnan:** Conceptualization, Methodology, Validation, Formal analysis, Investigation, Writing - Original Draft, Visualization, Writing - Review & Editing. **Blake Krisko:** Investigation, Visualization, Writing - Review & Editing. **Bulbul Ahmmed:** Investigation, Visualization, Writing - Review & Editing. **Luna Witchey:** Investigation, Writing - Review & Editing. **Oceani Valenzuela:** Investigation, Writing - Review & Editing. **Minas Mark Minasyan:** Investigation, Writing - Review & Editing. **Anthony Pak:** Investigation, Writing - Review & Editing. **Haik Mkhikian:** Conceptualization, Methodology, Validation, Formal analysis, Investigation, Writing - Original Draft, Visualization, Writing - Review & Editing, Supervision, Project administration, Funding acquisition.

## Data Availability

The mass spectrometry proteomics data have been deposited to the ProteomeXchange Consortium (http://proteomecentral.proteomexchange.org) via the PRIDE partner repository with the dataset identifier PXD068422"

## Supplementary Data

This article contains supplemental data.

## Declaration of generative AI and AI-assisted technologies

No AI or AI-assisted technologies were used in this work.

## Supplementary Figures

**Fig. S1.**
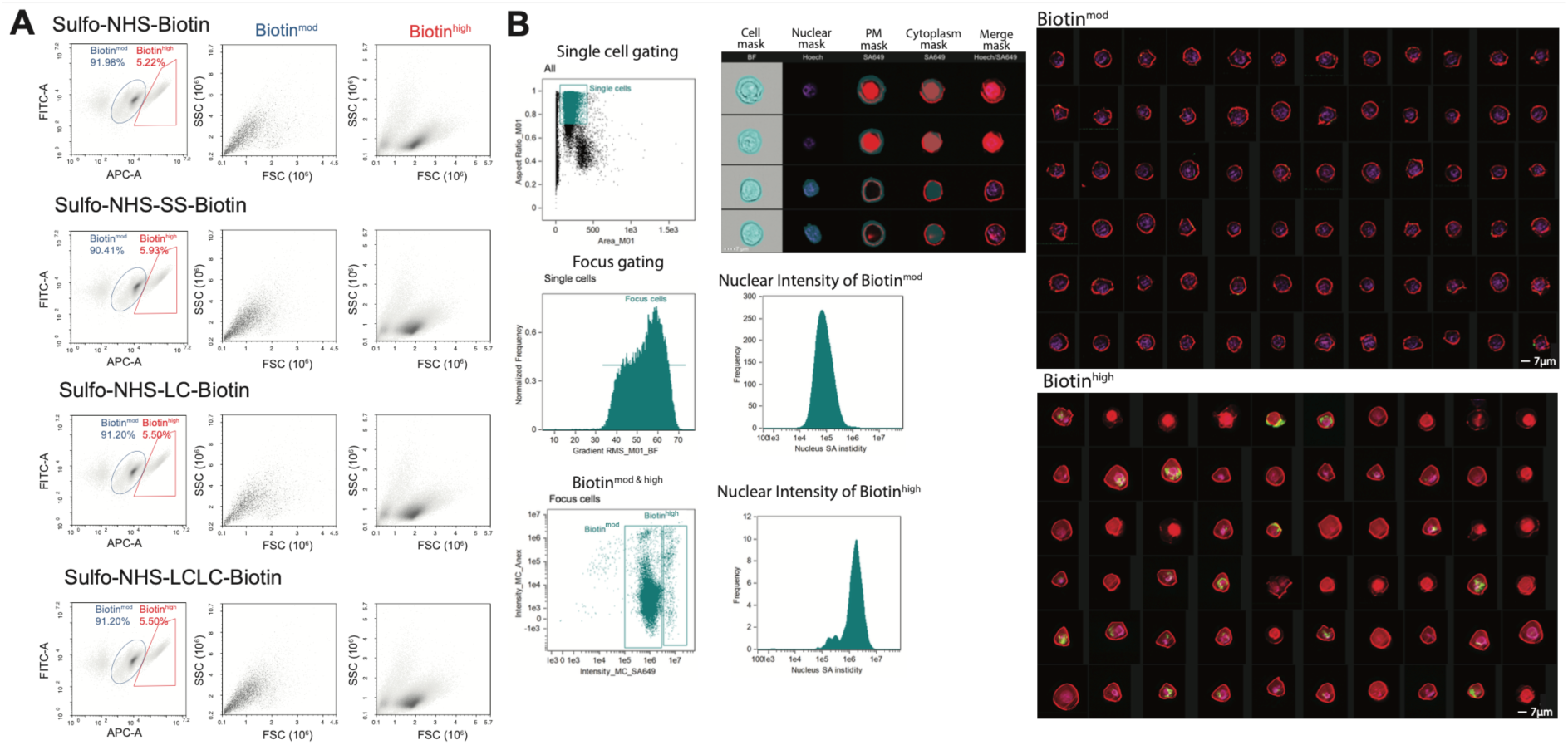
Evaluation of intracellular contamination by commonly used cell-impermeable biotinylation reagents. (A) Scatter plots illustrating two distinct populations Biotin^mid^ and Biotin^high^ following cell surface labeling with various cell-impermeable biotinylation reagents (left panels). The Biotin^high^ population (middle panels) exhibits reduced forward scatter (FSC) and increased side scatter (SSC), consistent with compromised membrane integrity and dead cell contamination. (B) Gating strategy for distinguishing Biotin^mid^ and Biotin^high^ populations and quantifying intracellular biotin distribution. Sulfo-NHS-biotin–labeled cells were stained with Annexin V–FITC, streptavidin-AF649 (SA-649), and DAPI, and 20,000 events per replicate were acquired by ImageStream®X Mark II imaging flow cytometer. Data were compensated using single-stained controls. Single cells were gated using brightfield (BF) area versus aspect ratio, and in-focus cells were selected by BF gradient RMS. A plasma membrane (PM) mask was generated by subtracting an eroded (0.7) BF mask from the original BF mask. Nuclear masks were defined using DAPI intensity to quantify biotin signal at the PM and in the nucleus.

**Fig. S2.**
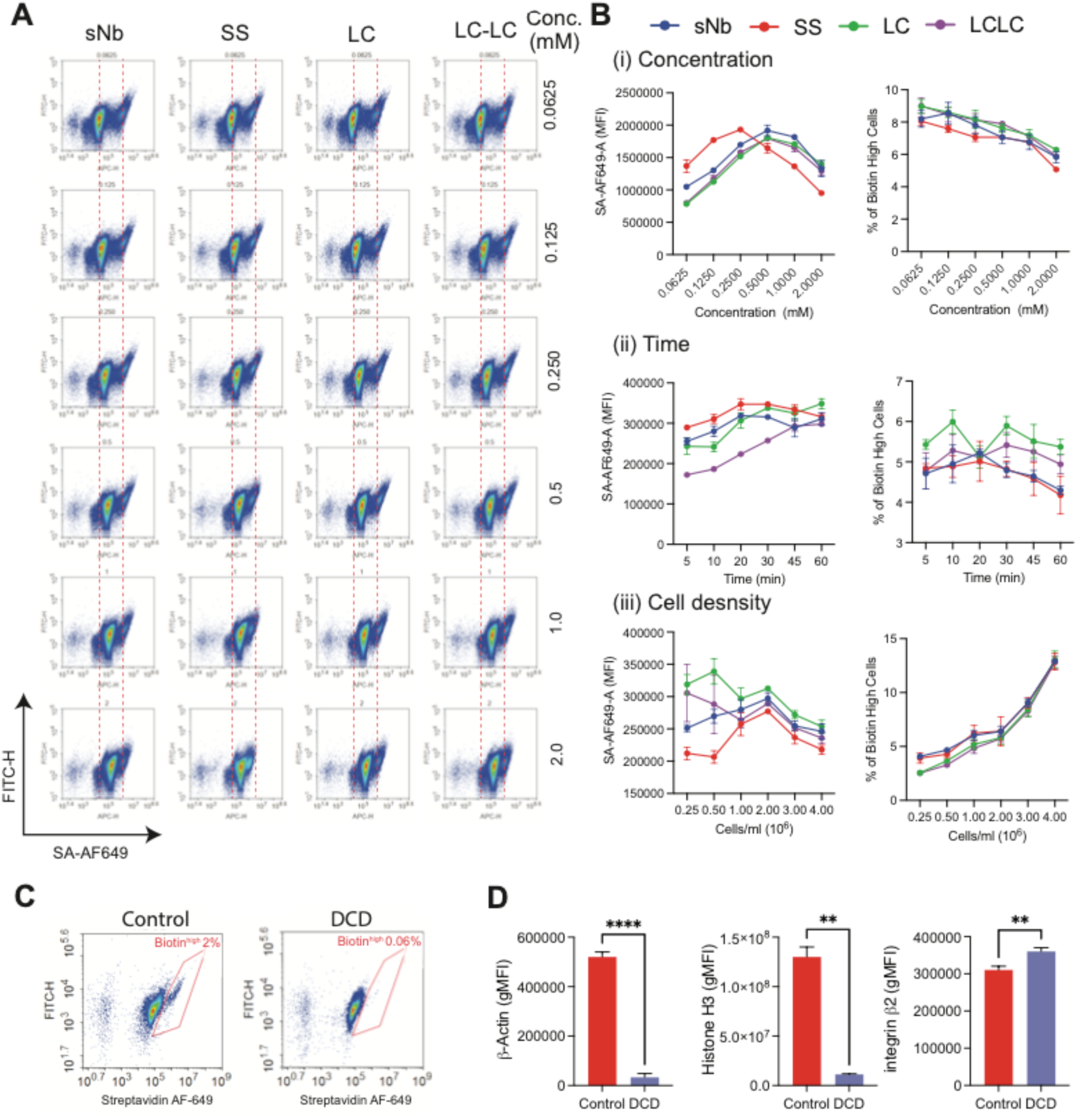
Optimization of biotinylation conditions to minimize intracellular contamination. (A) Representative scatter plots showing the effect of various biotinylation reagents on Biotin^mid^ and Biotin^high^ populations. Surface labeling was performed with sulfo-NHS-biotin (sNb), sulfo-NHS-SS-biotin (SS), sulfo-NHS-LC-biotin (LC), and sulfo-NHS-LCLC-biotin (LC-LC) as described. (B) Line graphs summarizing the effects of biotinylation reagent concentration, incubation time, and cell density on intracellular biotin signal (left panel) and the proportion of Biotin^high^ cells (right panel). For dose-response experiments, cells were labeled with increasing concentrations of each reagent. For time-course experiments, cells were incubated with 0.5 mM of each reagent for the indicated durations. To assess cell density effects, varying numbers of cells were labeled with 0.5 mM sulfo-NHS-biotin reagent for 20 minutes. Labeled cells were stained with streptavidin-AF649 and analyzed in triplicate by flow cytometry. (C) Representative scatter plots showing CD4⁺ T cells expanded with IL-7 and IL-15, with or without Annexin V–based dead cell depletion. Human CD4⁺ T cells were cultured in the presence of 5nM IL-7 and IL-15 for 14 days, then surface-labeled with sulfo-NHS-biotin, stained with streptavidin-AF649, and analyzed by flow cytometry. (D) Flow cytometric analysis of streptavidin-agarose bead-enriched biotinylated proteins from control and DCD samples. Cells were biotinylated with or without Annexin V-based dead cell depletion, lysed in 8 M urea, and enriched overnight with streptavidin-agarose beads. Enriched beads were stained with β-actin-FITC, Histone H3-PE, and Integrin β2-APC, and analyzed by flow cytometry. **p < 0.01; *p < 0.001 by unpaired two-tailed t-test with Welch’s correction.

**Fig. S3.**
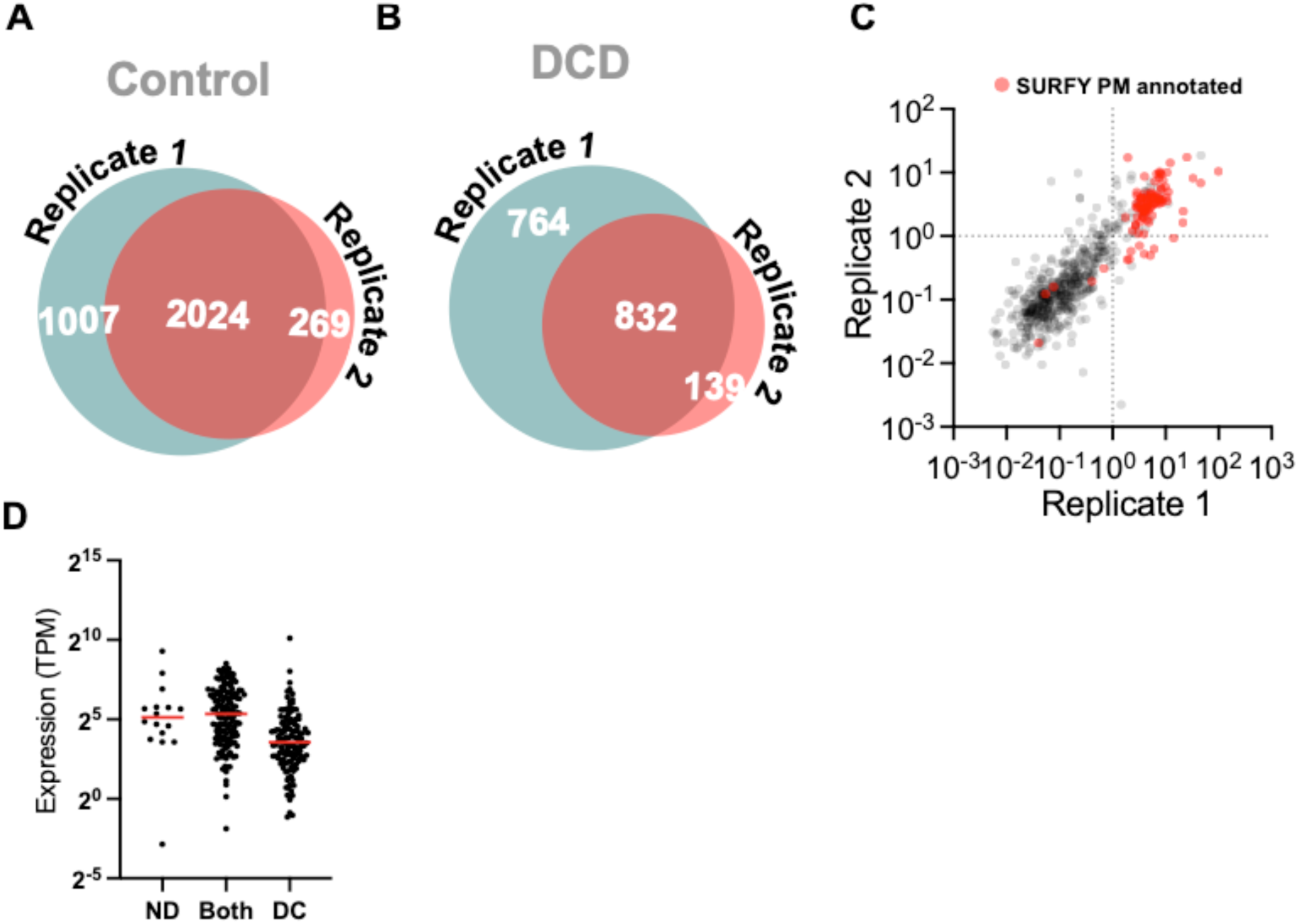
Reproducibility of dead cell depletion in plasma membrane protein detection. (A, B) Venn diagrams showing the overlap of quantified proteins across biological replicates in Control *(A)* and Annexin V-based dead cell-depleted (DCD, *B*) samples. (C) Scatter plot depicting the correlation of protein enrichment between DCD and Control samples. Non–plasma membrane (PM) proteins are shown in gray; SURFY-annotated PM proteins are shown in red. (D) Dot plot displaying mRNA expression levels of PM proteins identified exclusively in Control, exclusively in DCD, or shared between both. Jurkat transcriptome data were retrieved from the Human Protein Atlas.

## Supplementary Tables

**Table S1:** Proteomic and transcriptomic profiling of proteins identified in control and DCD samples.

**Table S2:** List of reference plasma membrane proteins and Topology annotation of identified proteins in Control and DCD samples.

**Table S3:** List of PM and Non-PM that are uniquely Identified in Control and DCD samples

**Table S4:** Annotation of Subcellular distribution and Comparison of reference PM annotation with enrichment score.

